# Cell composition inference and identification of layer-specific transcriptional profiles with POLARIS

**DOI:** 10.1101/2022.12.08.519631

**Authors:** Jiawen Chen, Tianyou Luo, Minzhi Jiang, Jiandong Liu, Gaorav P Gupta, Yun Li

**Affiliations:** Department of Biostatistics, University of North Carolina at Chapel Hill, Chapel Hill, North Carolina, USA; Department of Applied Physical Sciences, University of North Carolina at Chapel Hill, North Carolina, USA; Department of Pathology and Laboratory Medicine, University of North Carolina at Chapel Hill, Chapel Hill, North Carolina, USA; Department of Radiation Oncology, The University of North Carolina at Chapel Hill, Chapel Hill, NC 27599, USA; Lineberger Comprehensive Cancer Center, The University of North Carolina at Chapel Hill, Chapel Hill, NC 27599, USA; Department of Genetics, University of North Carolina at Chapel Hill, Chapel Hill, North Carolina, USA; Department of Computer Science, University of North Carolina at Chapel Hill, Chapel Hill, North Carolina, USA

**Keywords:** Spatial transcriptomics, deconvolution, differentially expressed genes, single cell, deep learning

## Abstract

Spatial transcriptomics (ST) technology, providing spatially resolved transcriptional profiles, facilitates advanced understanding of key biological processes related to health and disease. Sequencing-based ST technologies provide whole-transcriptome profiles, but are limited by the non-single cell level resolution. Lack of knowledge in the number of cells or cell type composition at each spot can lead to invalid downstream analysis, which is a critical issue recognized in ST data analysis. Methods developed, however, tend to under-utilize histological images, which conceptually provide important and complementary information including anatomical structure and distribution of cells. To fill in the gaps, we present POLARIS, a versatile ST analysis method that can perform cell type deconvolution, identify anatomical or functional layer-wise differentially expressed (LDE) genes and enable cell composition inference from histology images. Applied to four tissues, POLARIS demonstrates high deconvolution accuracy, accurately predicts cell composition solely from images, and identifies LDE genes that are biologically relevant and meaningful.

## Introduction

Molecular analysis of messenger RNA patterns in histological tissue sections is a key component of biomedical research and diagnostics. The development of novel spatial transcriptomic (ST) technologies has advanced dramatically over the last few years. There are two main categories of ST technologies: imaging-based or sequencing-based. Technologies based on imaging directly image individual RNA molecules within single cells [1, 2]. Sequencing-based techniques first label spatial spots on histological tissue sections with unique barcodes to indicate their two-dimensional spatial positions, and utilize RNA-sequencing to provide gene expression quantifications for each spot along with the spatial coordinates [3, 4]. Commonly used methods include MERFISH [1], seqFISH+ [5] in the former category and 10X Genomics’ Visium platform [3] in the latter category. More information can be found in recent review papers [6-8]. Some of the sequencing-based techniques (exemplary platforms include Spatial Transcriptomics and Visium) also provide a co-registered hematoxylin and eosin (H&E) stained histology image for the analyzed sample. Empowered by these technologies, we can obtain gene expression profiling with retained spatial information and histological images, which enable researchers and clinicians to gain a new level of insight into complex tissue samples.

In parallel to these technological developments, computational methods to analyze spatial data derived from tissue samples have substantially advanced. For instance, focusing on histology images, multiple machine learning and deep learning methods have been developed to maximally extract information from these images [9-12]. In the presence of pathological annotations, histology images can be used for various purposes including cell segmentation [13, 14], tissue type registration [15], mutation rate inference [16, 17], and gene expression prediction [12]. Most of these tasks, however, require pathologists to fully annotate each cell in the histological image, entailing substantial manual time and human resources. Such pathologist annotation is currently unavailable for the vast majority of publicly available ST data, thus making using traditional cell detection methods to perform ST deconvolution inaccessible. In the field of ST, histology imaging has primarily been used to predict gene expression and perform tissue registration, where the image data is usually subject to a pre-trained model to extract image features [9, 10, 12]. Several popular pre-trained models, such as convolutional neural networks, stacked sparse autoencoders, and masked autoencoders (MAE), have been employed as a first step to reduce image dimensions and demonstrate advantages in many applications [9, 10, 12, 13, 15]. However, cell composition inference hasn’t benefited from these models yet. In recent literature, histology images have been utilized to improve deconvolution accuracy [18, 19] but methods that can predict cell composition *solely* from histology images are currently unavailable.

Besides the histology image, ST data allow for the extraction and revelation of tissue structure through coordinated gene expression. Researchers have developed methods such as SPARK [20] and SpatialDE [21] for identifying genes whose expression varies within a tissue slice, known as spatially differentially expressed (SDE) genes. Gene expression changes spatially across spots within a tissue slice, often reflecting some underlying structured heterogeneity such as anatomical layers, clusters of similar spots, and/or spatial domains. Such structured heterogeneity motivates the development of ST clustering methods including BayesSpace to identify layers/clusters within each ST slice [11, 22, 23]. As aforementioned, the identified layers often correspond to different functions or morphological changes in the tissue [22, 24, 25]. The across-spot variation in expression can be largely attributed to three factors: variation in cell number, variation in cell composition, and true spatially driven variation in gene expression profile (**Fig. 1a**).

**Fig. 1.**
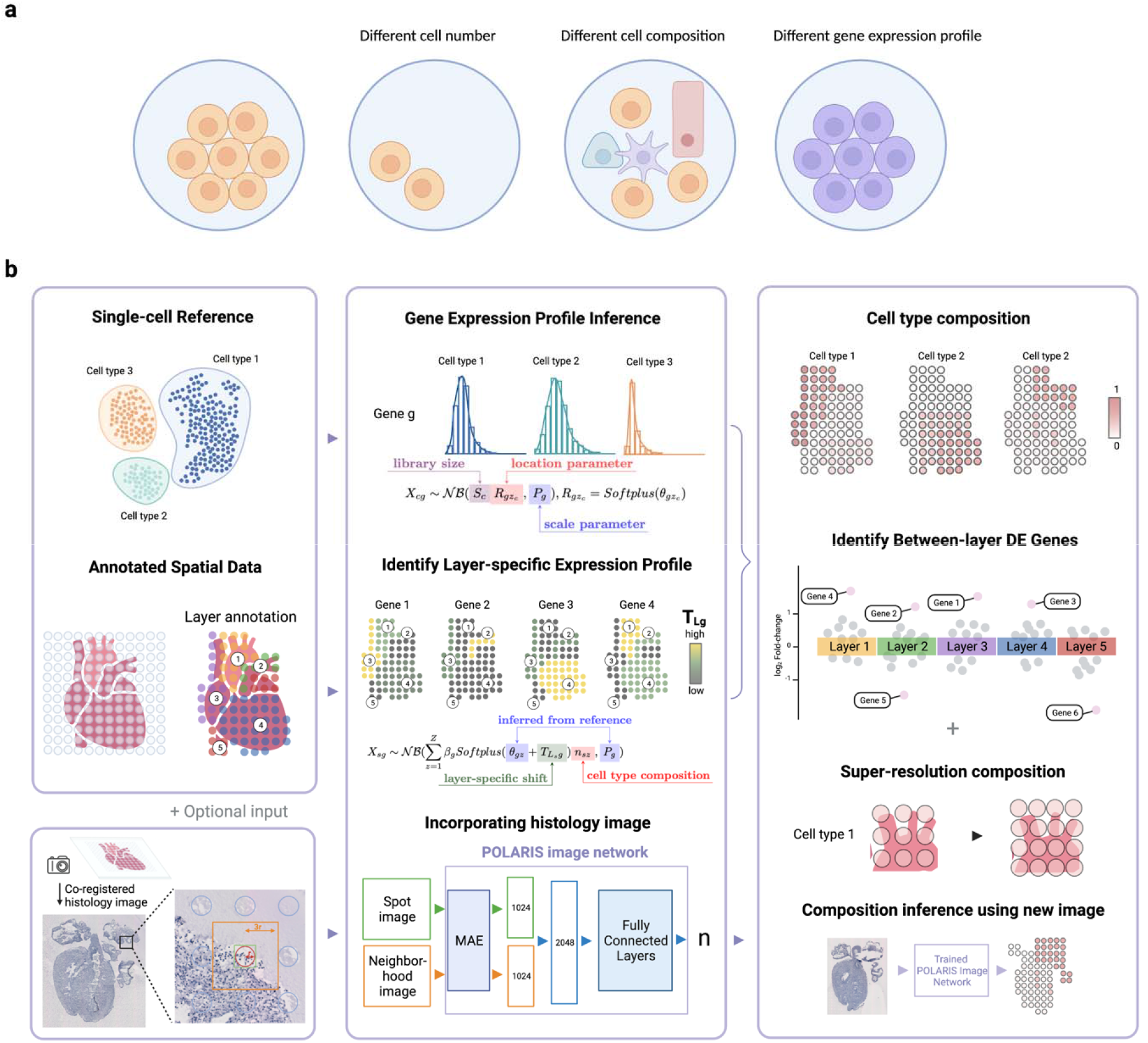
POLARIS overview. (a) Three reasons that can explain gene expression variation across spots. Each circle represents a spatial spot. Compared to the leftmost spot, the three spots on the right differ primarily in cell number, cell composition and gene expression profile. (b) POLARIS workflow. POLARIS takes as input single-cell reference and annotated spatial data, infers cell-type-specific gene expression profiles and identifies layer-specific expression profiles. When a co-registered histology image (as an optional input) is provided, POLARIS will additionally train an image network. The output of POLARIS includes inferred spot-level cell type composition, identified layer-wise differentially expressed (DE) genes and a pre-trained POLARIS image network that can be applied to independent images.

When the expression variation is truly driven by difference in spatial coordinates (in contrast to difference in cell number or composition), we can consider having sub-cell types located in different spatial regions (**Fig. 1a**). However, when SDE genes are detected by the aforementioned methods, the identified spatial difference is a result of the interplay of all three factors, and it is difficult to distinguish genes that are truly spatially differentially expressed from those that merely *appear* so due to differential cell number or composition across spatial spots. We would be able to differentiate among the driving factors if we had single-cell resolution data with the entire transcriptome or at least a large number of genes measured. However, in practice, we normally do not have this luxury: we have either data from imaging-based technologies that are single-cell resolution but measure only a small number of genes, or data from sequencing-based methods that provide transcriptome-wide measurement, but are limited in resolution. Sequencing-based ST methods have spatial spots of 2-100 *µm* in diameter, implying that each spot can easily contain tens of cells of different cell types. Lacking ideal (i.e., single-cell resolution with many genes measured) data motivates the development of computational methods to infer variation in gene expression profile across layers, while simultaneously estimating and adjusting for the estimated cell number and composition.

As a matter of fact, lack of knowledge in the number of cells at each spot nor the cell type composition itself has been recognized as a critical issue in ST data analysis, because failure to adjust for this accurately can lead to invalid downstream analyses. In order to address this problem, a number of ST deconvolution methods have been developed [18, 26-30]. However, most ST deconvolution methods assume that the gene expression profile for the same cell type is invariant across the entire tissue sample, which is a strong assumption whose violation will result in inaccurate cell composition inference. Methods such as DestVI that assume a continuous or smoothly changing gene expression profile across the tissue, however, have exhibited inconsistent performance across tissue types [6, 31]. Therefore, how to model layer-specific gene expression variation and utilize histological images to infer cell composition is a problem that remains unsolved.

Here we present POLARIS, Probabilistic-based cell cOmposition inference with LAyer infoRmatIon Strategy, to perform cell type deconvolution and infer layer-wise differentially expressed (LDE) genes (**Fig. 1b**). POLARIS integrates single-cell RNA-seq reference and ST data with annotated layer information. By examining histology images and the coordinated expression profile, one can reasonably infer layers or sub-regions that correspond to different biological functions (e.g. cancer vs non-cancer regions in a tumor biopsy, different layers in a brain cortical sample, ventricle and atrium areas in heart). By explicitly allowing and modeling layer-specific gene expression patterns, POLARIS is not only capable of identifying cell type composition with high accuracy, but also could identify LDE genes while simultaneously correcting for differential cell composition. An additional key characteristic of POLARIS is its flexibility to optionally leverage histology images. To our knowledge, POLARIS is the first ST deconvolution method that can predict cell composition purely from a histological image. This functionality also empowers POLARIS to infer super-resolution cell composition based on images of areas without gene expression measurements (i.e., areas in between spots), as well as to predict cellular composition based purely on an original H&E stained image. The performance of POLARIS was evaluated on data from multiple tissues including the mouse cortex, developing human heart, and HER2+ breast cancer samples. POLARIS robustly demonstrates high deconvolution accuracy across tissues compared to other state-of-the-art deconvolution methods, accurately predicts cell composition solely from images, and identifies LDE genes that are biologically relevant and meaningful. Our results showcase the advantages of POLARIS in the following three aspects: deconvolution accuracy, LDE gene identification and prediction with image.

## Result

### POLARIS method overview

POLARIS is a probabilistic-based inference method that assumes that gene expression counts in both scRNA-seq reference data and ST data follow a negative binomial distribution. As a first step, POLARIS maximizes likelihood to infer cell-type-specific gene expression profiles from scRNA-seq reference (**Fig. 1b**). The gene expression profile of each spot in ST data can then be viewed as a weighted sum of the negative binomial distribution derived from the scRNA-seq reference, where the weights are based on spot-level cell composition. As opposed to assuming that cell-type-specific gene expression profiles are invariant throughout a whole tissue slice, POLARIS assumes that only spots in similar biological or anatomical layers share the same gene expression profiles by introducing a layer-specific location parameter. Explicitly modeling layers is a unique feature of our POLARIS method. POLARIS accepts any user-specified layer annotations, e.g., derived manually (from pathologist annotation) or computationally (based on either morphological features or gene expression, e.g., using BayesSpace [22]). Note that the layer-specific parameters cannot be inferred from single-cell reference because there’s no layer information by the nature of data generation. Using ST data with layer annotations, POLARIS enables layer-specific inference. By introducing the layer-specific shift parameters (**Methods**), we can obtain an updated location parameter for each layer in the ST data, allowing cell-type-specific gene expression profiles to vary across layers. By multiplying the updated location parameter with the cell composition parameter as well as the parameter to account for technical/batch effects, we simultaneously model the impact of cell composition and spatial location (as reflected by layers) on cell-type-specific gene expression, while controlling for potential batch effects. Parameters can be estimated using maximum a posteriori estimation (MAP) (**Methods**). So far, we have focused on inference with gene expression data only. When a co-registered histology image is available, POLARIS first employs MAE [32] to extract features from the image tile of each spot and the image tile of its neighborhood. These two extracted features are then combined and used as inputs to build POLARIS’s image network (**Fig. 1b**). The output of POLARIS’s image network is cell composition for any input image (which can be from a completely independent histological image). The output of POLARIS includes inferred spot-level cell type composition and layer-specific gene expression profiles, as well as a trained POLARIS image network. The layer-specific gene expression profiles enable identification of LDE genes, and the pre-trained POLARIS image network allows resolution enhancement and cell composition inference from a new histology image.

### POLARIS attains high deconvolution accuracy

The deconvolution accuracy of POLARIS was assessed both through simulation and in single-cell resolution ST data. Specifically, we simulated cells with gene expression counts from cell-type and layer-specific negative binomial distributions and randomly selected cells to create spot-level gene expression. For single-cell resolution real ST datasets, we clumped cells into spots according to their coordinates to mimic low-resolution spot-level ST data. We used data where we have cell type labels for the single cells such that we have the true cell type mixture in each clumped pseudo-spot. We quantified the performance using root mean square error (RMSE) where a smaller RMSE corresponds to better performance. We compared POLARIS with five state-of-art methods: CARD [26], DestVI [29], RCTD [28], stereoscope [27] and SPOTlight [30].

We began with a simulated scenario where all spots and layers share a similar composition of cells, but with layers differing in terms of their gene expression profiles. Under this scenario, gene expression variations are solely the result of variations in gene expression profiles across layers. Specifically, we first simulated a dataset with two “biological” layers, with cells from six cell types and expression values for 100 genes generated. We first simulated the layer of each cell and then the gene expression values for the cell were drawn from negative binomial distributions according to its layer and cell type. We then constructed pseudo spots by randomly selecting 10-16 cells from each layer. Specifically, we generated 200 spots with 50 spots in layer 1 and 150 spots in layer 2 (**Methods**). Under this simulation framework, genes could be classified into three categories: up-regulated in layer1 (e.g., *Gene36*), up-regulated in layer2 (e.g., *Gene100*) and no significant difference between layers (e.g., *Gene95*) (**Fig. 2a-b**). Applied to the simulated data, POLARIS outperforms all other methods as manifested by its lowest RMSE (**Fig. 2c**).

**Fig. 2.**
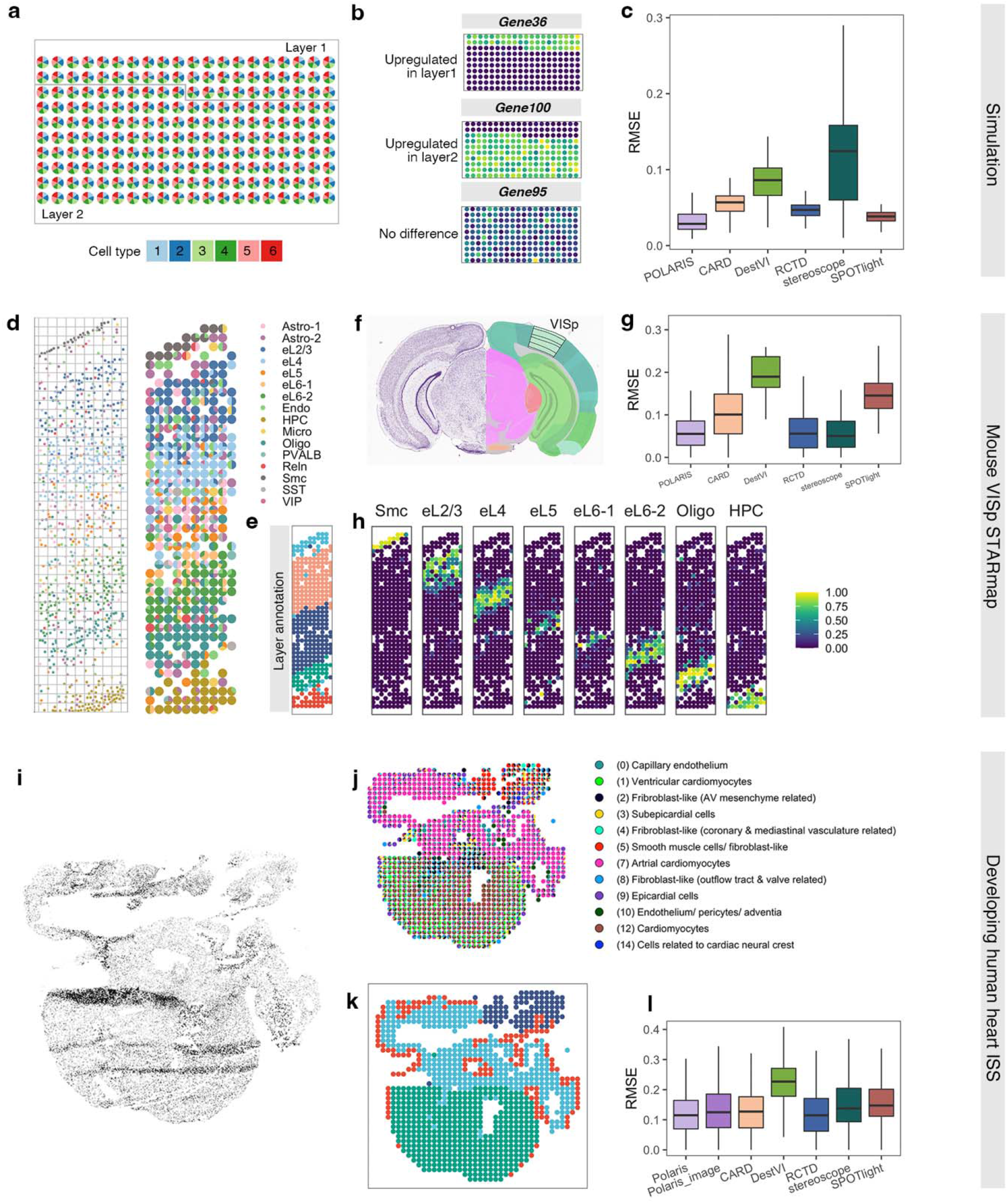
Deconvolution accuracy of POLARIS. Simulation: (a) cell type composition of the simulated ST data. (b) Three categories of gene expression pattern: up-regulated in layer1, up-regulated in layer2, no significant difference across layers. (c) RMSE of POLARIS along with other state-of-the-art methods on simulated data. **On mouse VISp STARmap data**: (d) (left) cell map and (right) clumped pseudo spots along with their cell compositions visualized by pie charts (e) BayesSpace identified clusters, consistent with layer structure of mouse VISp (f) Nissl staining (left) and anatomical annotations (right) from the Allen Mouse Brain Atlas and Allen Reference Atlas - Mouse Brain. The black lined area indicates the layer structure of the VISp region. (g) RMSE of POLARIS along with other state-of-the-art methods on mouse VISp STARmap data. (h) POLARIS inferred spot-level composition of the eight major cell types, along cortex depth. **On developing human heart ISS data**: (i) Processed DAPI-stained histology image of the developing human heart ISS data. (j) clumped pseudo spots in the ISS data along with their cell compositions visualized by pie charts. (k) BayesSpace identified clusters in the ISS data. (l) RMSE of POLARIS along with RMSE of other state-of-the-art methods on the heart ISS data.

We further assessed POLARIS’s deconvolution performance in a real single-cell resolution ST data from the mouse VISp region, a well-structured region in mouse cortex that has been extensively studied [2, 33]. Anatomical structure, major cell types, and layer-specific gene markers provide information about the layered and segmented structure of the mouse VISp (**Fig. 2d-f**) [33, 34]. We used single-cell resolution ST data from the STARmap platform [2], which consists of 1,020 genes measured in 973 cells. We divided the cells into 356 pseudo spots each of 400×400 square pixels (**Fig. 2d**). In order to perform deconvolution, we utilized the internal reference (that is, the STARmap single cell data itself as the reference). In this way, any systematic differences between the reference and the target ST data are eliminated as potential factors that may impair performance. This internal reference evaluation provides a baseline (or upper bound) for measuring the performance of deconvolution methods [6]. Since layer annotation is required when using POLARIS, we employed BayesSpace [22] to cluster the constructed pseudo spots, resulting in five distinct clusters, reflecting the expected layer structure of mouse VISp (**Fig. 2e**). In this mouse VISp dataset, POLARIS still achieves amongst the best performance in terms of RMSE (**Fig. 2g**). Moreover, POLARIS, based on its inferred cellular composition, successfully recovers the layer structure of mouse VISp (from top to bottom: Smc, eL2, eL3, eL4, eL5, eL6-1, eL6-2, Oligo, HPC, **Fig. 2h**).

Additionally, we tested POLARIS on the developing human heart tissue generated from the in situ sequencing (ISS) platform (**Fig. 2i**) [35]. The heart ISS data is also a single-cell resolution ST data, consisting of 24,371 cells and with only 65 genes measured in each cell. We gridded the cells into pseudo spots each of dimension 454 × 424 square pixels (**Fig. 2j**). The main purpose of this assessment is to evaluate POLARIS’s performance with a limited number of genes. Instead of using the internal reference (ISS data itself), we used a scRNA-seq reference obtained from a similar biological sample [36]. Consequently, we can also evaluate the deconvolution performance when the reference and ST data are not perfectly matched. The heart ISS data provides us with a DAPI stained histology image, allowing us to measure the performance of POLARIS by including the histology image as an additional input (**Fig. 2i, Methods**). We clustered the spots into four layers using BayesSpace (**Fig. 2k**). The BayesSpace inferred layers correspond reasonably well to the anatomy of the heart (red: epicardium, green: ventricles, light blue: atria, dark blue: outflow tract). POLARIS has maintained its best performer position. Specifically, POLARIS achieves the lowest/best mean of MSE (**Fig.2l**). In spite of the fact that the DAPI staining only contains one color channel, POLARIS with image input is able to effectively infer the type of cell, achieving accuracy close to the best performers.

### Polaris identifies layer-specific gene expression pattern

A major feature of POLARIS is its ability to model layer-specific parameters. The layer/structure of a tissue can be reflected in multiple dimensions, such as morphology, gene expression, and other omics levels. POLARIS focuses on leveraging the rich gene expression information provided by ST data. As detailed above, cell density, cellular composition, and the “real” spatially differentially expressed genes can all contribute to the observed gene expression variation. By incorporating layer-specific parameters into the cell type deconvolution process, POLARIS is able to identify such LDE genes while taking into account differential cell composition. POLARIS quantifies statistical significance for LDE genes using permutation tests, and magnitude of effect using log2 fold change in mean gene expression, based on the inferred layer-specific mean parameters (**Methods**). Through the elimination of potential confounding effects of cell composition, POLARIS ensures that the LDE genes identified are differentially expressed genes truly due to spatial factors.

As a starting point for assessing POLARIS’ ability to infer LDE genes, we performed simulations where we know the truth. Following the same simulation framework used above to evaluate deconvolution efficiency, we evaluated the layer-specific location parameters. Again, since cellular composition is simulated from the same distribution across spots regardless of layer status, observed gene expression variation can only be attributed to truly differential expression patterns across layers (**Fig. 2a-b**). Consequently, genes could be classified into three categories: layer1-enriching genes, layer2-enriching genes, and genes with similar expression levels across layers (**Fig. 2b, 3a**). We applied POLARIS to perform deconvolution and simultaneously perform the permutation test and calculate the log2 fold change in mean expression, layer1 over layer2. POLARIS successfully identified genes that have different expression profiles across layers (**Fig. 3b**). The predicted log2 fold change well captures the true log2 fold change (**Fig. 3c**) when all the genes have layer-specific gene expression profiles. In this particular simulation, we generated the genes such that all of them have layer-specific expression profiles, although the expression difference between layers of some genes could be small (detailed in **Methods**). To investigate the impact of the proportion of LDE genes on POLARIS performance, we further conducted simulations using the same setting but varying the proportion of genes with layer-specific expression profiles (**Extended Data Fig. 1**). As the proportion of LDE genes increases from 0 to 1, UMAP representations of cells become more separated by layers. It is difficult, however, to identify the sub-types of cells based on the UMAP representation, even with prior knowledge of the layer information when the proportion of LDE genes is < 0.5, thus failing to capture heterogeneity across layers (**Extended Data Fig. 1a**). POLARIS well controlled the type-I error and accurately estimated the true log2 fold change regardless of the proportion of LDE genes (**Extended Data Fig. 1b**). Note that layer-specific parameters are inferred only from the ST data because we do not have layer information from the scRNA-seq reference. As a result, the realistic data we have is not ideal for layer-specific inference. POLARIS, nevertheless, is able to reveal the LDE genes.

**Fig. 3.**
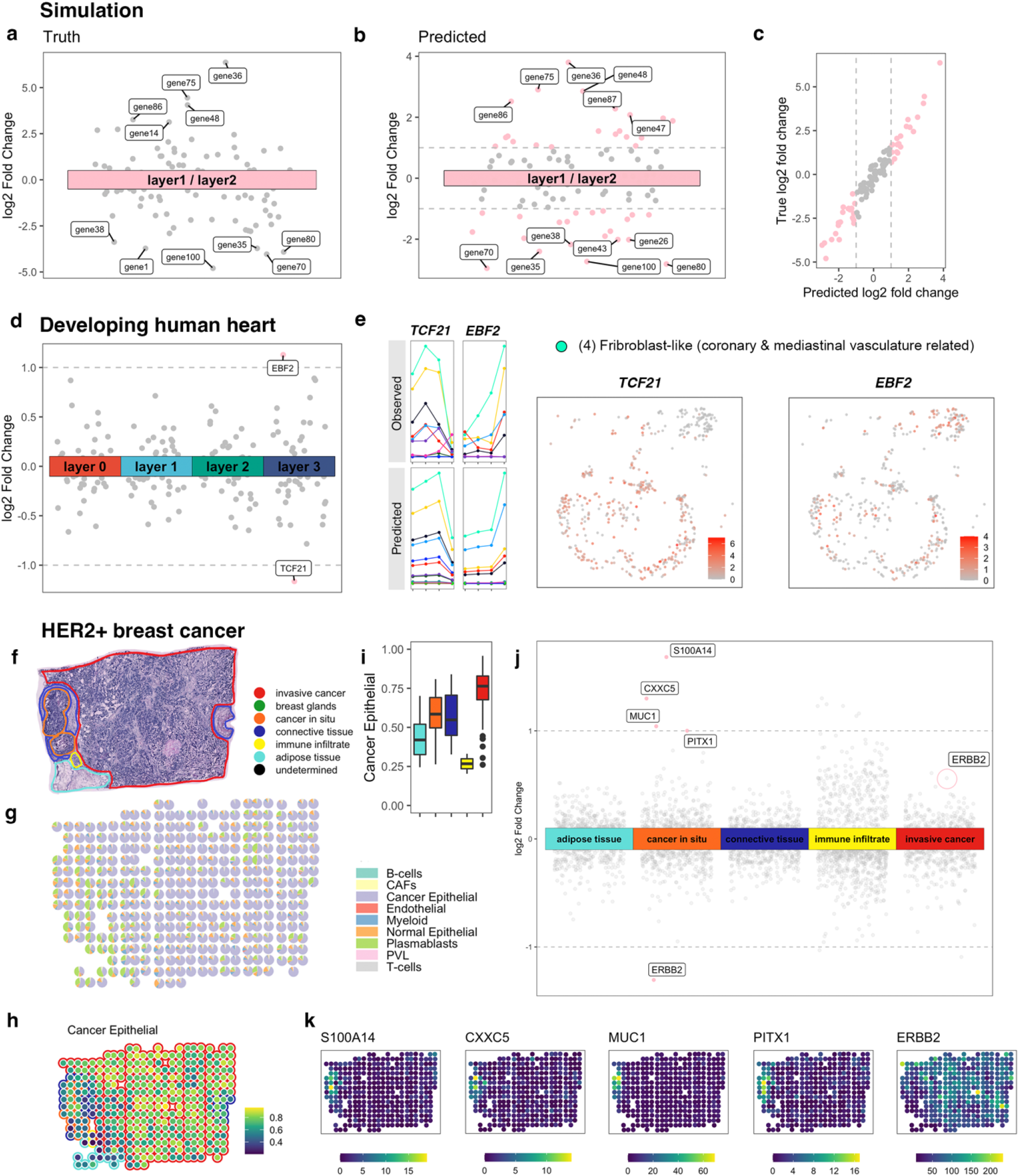
LDE genes identified by POLARIS on three datasets. On simulated data: (a) true log2 fold change of mean gene expression, layer1 over layer2. (b) POLARIS inferred log2 fold change of mean gene expression, layer1 over layer2. POLARIS identified LDE genes are colored as pink, otherwise, gray. (c) POLARIS inferred log2 fold change of gene expression across layers achieves high correlation (Pearson’s correlation = 0.974) with the true log2 fold change. **On developing human heart data**: (d) POLARIS inferred log2 fold change of gene expression across layers. POLARIS identified LDE genes are colored as pink, otherwise, gray. (e) (left) Observed mean gene expression in scRNA-seq data (top) and POLARIS inferred gene expression location parameter (bottom) of each cell type across layers. Lines are colored by cell types. X axis indicates layer status: from left to right is layer 0,1,2,3. (right) Gene expression of TCF21 and EBF2 in the Fibroblast-like (coronary & mediastinal vasculature related) cells. **On HER2+ breast cancer data**: (f) Pathologist annotation on slide A1. (g) POLARIS inferred cell composition (h) POLARIS inferred cancer epithelial cell proportion (i) Distribution of POLARIS inferred cancer epithelial cell proportions in each layer (color scheme is the same as in Fig. 3e) (j) POLARIS inferred log2 fold change of gene expression across layers. Points with absolute value greater than 1 are colored as pink, otherwise, gray. (k) Gene expression profiles of POLARIS identified LDE genes in the cancer in situ layer.

Encouraged by POLARIS’s performance in simulated data, we proceeded to further test POLARIS’s capability to detect LDE genes using real datasets. We began with the developing human heart single-cell resolution ISS ST data. We created pseudo-spots from the dataset and applied POLARIS to the pseudo-spots, feeding as input spot-level gene expression information along with layer information inferred from BayesSpace. LDE genes detected by POLARIS indeed have differential patterns of gene expression in different layers (**Fig. 3d-e)**. Although cell type affects gene expression, layer status also plays a significant role in shaping the spot-level expression profile. Furthermore, such layer-specific changes are shared by many cell types. For example, the expression of *EBF2* in most cell types is highest in layer 3 compared to the other three layers (**Fig. 3e**). POLARIS, by making accurate inference of gene expression profiles in different cell types, has successfully captured the LDE gene and has recovered the layer-specific variation present in most cell types. We further explore the links between LDE genes and biological functions of each layer. For example, among the LDE genes, *TCF21* has the lowest mean gene expression in layer 3 (**Fig. 3d-e**) where fibroblast-like cells are enriched (**Fig. 2j-k**). The transcription factor 21 (TCF21) encoded by *TCF21* plays a crucial role in regulating cell differentiation and cell fate determination through epithelial-mesenchymal transformations during cardiac development. More specifically, TCF21 has been reported to be capable of promoting the development of cardiac fibroblasts and inhibiting differentiation of epicardial cells into vascular smooth muscle cells [37], consistent with our observed down-regulation in layer 3.

We further applied POLARIS on a more complex tissue: breast cancer samples. There are several subtypes of breast cancer, among which the HER2-positive subtype is characterized by the increased expression of *ERBB2* (AKA *HER2*, human epidermal growth factor receptor 2) in tumor cells [38]. We obtained ST data of HER2-positive tumors from eight individuals (patient A-H) generated through the Spatial Transcriptomics platform [3, 38]. Each of the eight patients provided multiple slides, but only one slide from each patient was pathologist annotated. Annotations mark areas with one of the following five labels: in situ cancer (noninvasive ductal carcinoma in situ, DCIS), invasive breast cancer (IBC), adipose tissue, immune infiltrate, or connective tissue [38]. Here we highlight the results of two slides with both IBC and DCIS regions (results from the A1 slide in **Fig. 3f-k**, results from the G2 slide in **Extended Data Fig. 2**). We applied POLARIS on the pathologist annotated ST data using an external scRNA-seq reference [39]. POLARIS made reasonable inference regarding cell composition on slide A1 (**Fig. 3f**). For example, cancer epithelial cells are inferred to be enriched in the DCIS and IBC areas. POLARIS also identified several LDE genes in the DCIS area (**Fig. 3i-j**). These LDE genes, including *S100A14, MUC1, PITX1*, and *ERBB2* are mainly expressed in cancer epithelial cells (**Extended Data Fig. 3**). In general, genes that are primarily expressed in cancer epithelial cells are enriched either in the DCIS or the IBC region (**Fig. 3h-k**), which is expected since these two regions have similarly high proportions of cancer epithelial cells. The LDE genes identified by POLARIS successfully capture cancer epithelial cell specific genes, but differentially expressed in the two regions (**Fig. 3h-k**). For example, in the DCIS region, all POLARIS-identified LDE genes except *ERBB2* have a positive log2 fold change estimate (**Fig. 3j**), consistent with expression patterns shown in **Fig. 3k**. The observed down-regulation of *ERBB2* in the DCIS area most likely reflects an up-regulation of *ERBB2* elsewhere. As shown in **Fig. 3f**, the majority of slide A1 is the invasive cancer area. Cancer epithelial cells in the invasive cancer area presumably invade other areas resulting in an increased *ERBB2* expression in all other pathologist-identified areas except the DCIS and these invaded areas also exhibit an increase in the proportion of cancer epithelial cells (**Fig. 3h-i, Extended Data Fig. 2f**). ERBB2, the protein encoded by *ERBB2*, plays an important role in breast cancer. The over-expression of *ERBB2* disrupts normal cell-control mechanisms and gives rise of aggressive tumor cells and leads to increased breast cancer metastasis [40-45]. Interestingly, when applied to slide G2, *ERBB2* shows to be up-regulated in the DCIS area (**Extended Data Fig. 2e,g**). Together, these observations reflect the heterogeneity of the samples and are consistent with the literature that *ERBB2* is over-expressed in 30-35% of DCIS, while *ERBB2* is only expressed in 15-25% of IBC [46-49]. POLARIS reveals such heterogeneity and complexity by showing differential gene expression profiles between DCIS and IBC regions on the same slide, and by revealing differential patterns across slides from the same patient as well as across patients. In addition to *ERBB2*, other LDE genes identified by POLARIS and the proteins encoded by those genes also play important roles in breast cancer. For example, copy number amplification of *S100A14*, significantly correlated with the increased *S100A14* mRNA expression, is present in 5.4%-20.7% of primary breast cancer patients and in approximately 26.1% of metastatic breast cancer patients [50]. For another example, *CXXC5* over-expression has been observed to be associated with a poor prognosis for estrogen receptor positive (ER+) breast cancer [51].

### Polaris enables prediction purely from histology image

After demonstrating that even a single-color histology image is able to generate high-accuracy deconvolution inference in the developing human heart tissue, we continued to apply POLARIS to other spot-level ST data with H&E staining images. We utilized the mouse primary somatosensory cortex area (SSp) data generated from the 10x Visium platform [52]. Similar to the mouse VISp region, mouse SSp is also an area in the mouse cortex with well-defined anatomical and functional structure. Specifically, the glutamatergic neuron types exhibit clear layered patterns [6, 33, 53]. Using an independent scRNA-seq from similar SSp regions [33] as the external reference, POLARIS trained an image network using four SSp ST slides. Each slide was clustered into six groups using BayesSpace (**Extended Data Fig. 4**). POLARIS successfully captures expected patterns of glutamatergic composition and reveals layer structure consistent with data from the Allen Brain Atlas [34].

Spot-level ST technologies cannot measure every part of a tissue slide. As shown in **Fig. 4a**, gene expression levels are not available for any region outside the measured spots (**Fig. 4a**). POLARIS, through its trained image network, can determine cell type composition using the image of unmeasured areas. Specifically, POLARIS-trained image network can be applied to cropped images of the same size as the original grid in training. Sliding across the entire histological image of the SSp slide and applying the POLARIS-trained image network to the image of each sliding window, one can obtain super-resolution inference, encompassing areas not initially covered by spatial spots. Such super-resolution inference empowers us to gain finer details of the layered structure (**Fig. 4b**). Following the training of POLARIS’s image network, we applied it to a new SSp slide to test whether the trained image network is able to make reasonable inference on images from independent slides (**Fig. 4c**). We compared cell compositions inferred using the image network trained by POLARIS from other slides, with those inferred using the new slide’s own gene expression and histology image on measured spots (**Fig. 4d-e, Extended Data Fig. 5a-b**). Although the two sets of inference show differences, results obtained solely from POLARIS’s pre-trained image network still show strong correlation with the inference results using its own image and gene expression, especially in glutamatergic neurons (For example, the Pearson’s correlation for L6b CTX is 0.8). POLARIS-trained image network successfully recovers the layered structure in most cell types, suggesting that the approach could be applied to new histology slides as long as we have a network pre-trained by POLARIS using ST data from similar regions.

**Fig. 4.**
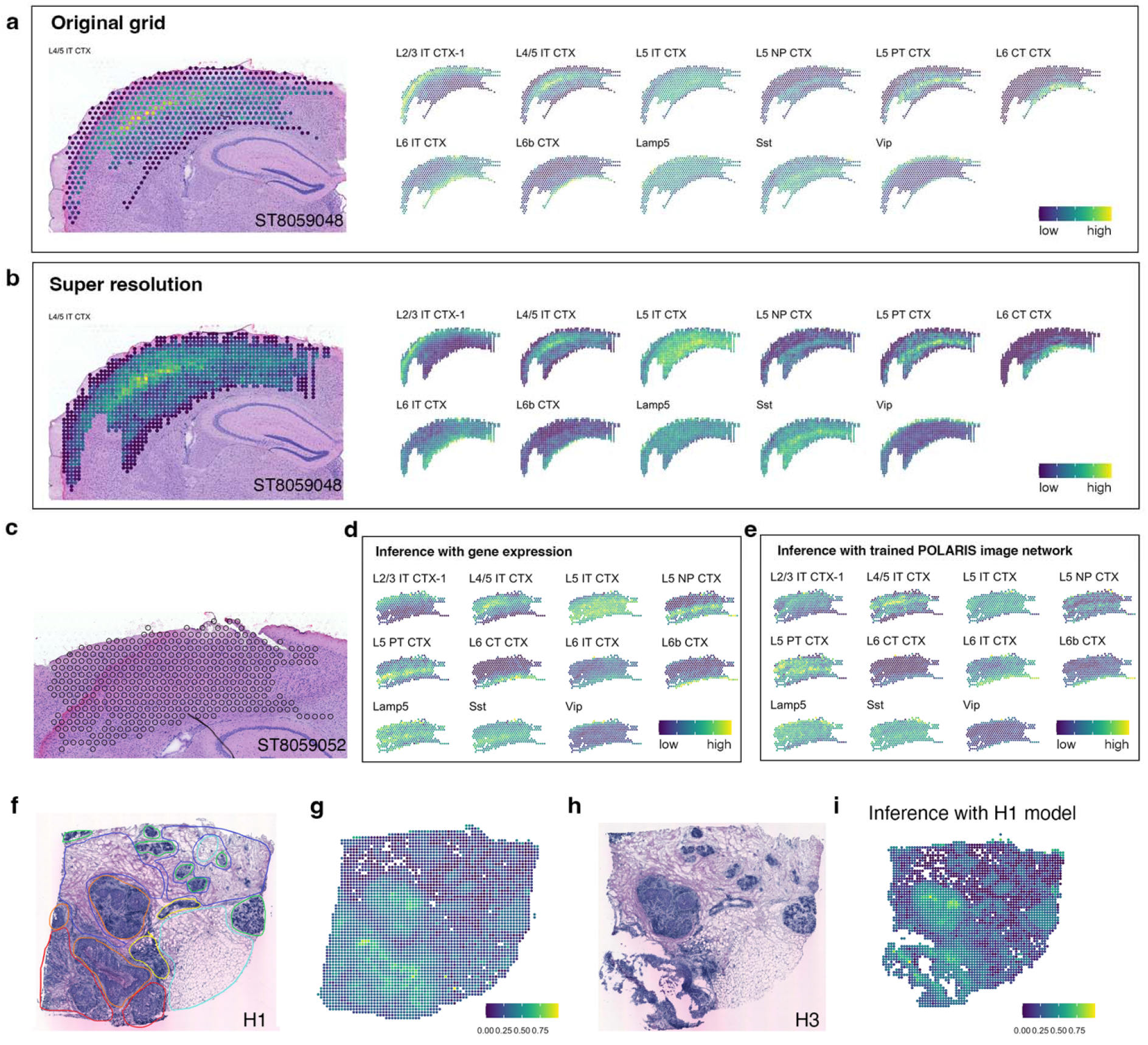
POLARIS achieves super-resolution cell composition inference when using histology image as input. On mouse SSp 10x Visium slide ST8059048: (a) The original grid and POLARIS inferred cell composition. (b) The super-resolution cell composition inferred using a POLARIS-trained image network. Points in (a) and (b) are colored by the corresponding cell proportion (from blue to yellow corresponds to low to high). **On slide ST8059052**: (c) The original grid of the slide (d) POLARIS inferred cell composition inference using spot-level gene expression and histology image. (e) Inferred cell composition using a POLARIS-trained image network (trained on the other 4 slides), based on histology image only. Similarly, points in (d) and (e) are colored by the corresponding cell proportion (from blue to yellow corresponds to low to high). **On Her2+ breast cancer**: (f) pathologist’s annotation of slide H1. (g) Super-resolution inference of H1 using POLARIS image network trained on H1. (h) Histology image of slide H3. (i) Super-resolution inference of H3 using POLARIS image network, again trained on H1. The points in (g) and (i) are colored by the cancer epithelial cell proportion. CTX: isocortex, IT: intratelencephalic, PT: pyramidal tract, NP: near-projecting.

POLARIS-trained image network was further examined using breast cancer data. We again used pathologist annotated ST data and the external scRNA-seq from Wu et al. as reference to perform deconvolution on slide H1 (**Fig. 4f**) [38, 39]. Afterwards, we employed the image network trained by POLARIS from slide H1 to enhance resolution and obtain super-resolution cell composition inference on the H1 slide itself (**Fig. 4g**). The inferred cell compositions are consistent with those in other slides where cancer epithelial cells are enriched in the DCIS and IBC regions. It appears that cancer epithelial cell proportion is able to accurately capture the cancerous areas. For example, spots with >0.4 proportion of cancer epithelial cells are enriched within regions labeled as DCIS and IBC by pathologists (**Extended Data Fig. 5e**). We then further applied the POLARIS image network trained on H1 to two other slides of the same patients (H2 and H3), which have no pathologist’s annotation (**Fig. 4h-i, Extended Data Fig. 5c-g**). By closely examining the histology images, it is evident that regions characterized by high proportions of cancer epithelial cells are indeed primarily cancerous areas. Our results therefore suggest that POLARIS enables a new method of registering histology images to different anatomical/functional regions, for example cancerous areas in this analysis. By examining cell composition in each spot, we are able to group the spots into layers. In summary, with a trained POLARIS image network, we could obtain super-resolution cell composition inference which reveals finer layer structure of a tissue.

POLARIS allows cell composition inference on new histology images without gene expression and consequently, is able to identify anatomical and functional regions. Compared to existing supervised classification methods for registering histology image tiles to different regions or layers which require annotated layer information, POLARIS is unsupervised in terms of histology information and does not require pathologist annotation. Instead, POLARIS relies on pre-knowledge about the relationship between the targeted layer and the cell composition.

## Discussion

ST technologies are rapidly evolving. In the near future, we expect to be able to measure gene expression levels at single cell resolution and of all genes in the transcriptome. Currently, spot-level resolution ST technologies such as the Visium and Spatial Transcriptomes still have their advantages in terms of throughput (both in terms of number of genes measured and number of spatial spots examined) and the ability to obtain high-resolution H&E staining images. Because of their advantages, researchers are generating a deluge of these data. It is, however, imperative to perform cell type deconvolution at each spot in order to mitigate or eliminate potential confounding caused by differential cell composition across spots. Despite numerous methods developed for ST deconvolution, two pieces of information have been under-utilized. First, often there is a layer structure, or at least areas reflecting different anatomical or functional regions in an ST slide. Second, histological images, carrying information complementary to spot-level gene expression profiles, have not been fully explored in their value for cell type decomposition. In this work, we present POLARIS, a unified framework that leverages layer structure information and/or histological images, for cell type deconvolution both at spots with expression measurement and in regions with only image information, as well as for the revelation of LDE genes.

We demonstrate the performance of POLARIS on simulation and real datasets including developing human heart, mouse cortex VISp and SSp region, and human HER2+ breast cancer samples. POLARIS robustly achieves best or close to best deconvolution performance compared to other state-of-the-art methods. POLARIS’s inference on spot-level ST data reveals layered structures that are consistent with gene expression profiles, histological images, and known/established anatomical/functional layers/regions in the corresponding tissue samples.

Equally if not more importantly, POLARIS accurately infers layer-specific expression profiles across different cell types which leads to the identification of LDE genes. We demonstrate POLARIS’s power to identify LDE genes using simulation data as well as the single-cell resolution ST data from the developing human heart, where we have knowledge regarding the true LDE genes. The LDE genes identified by POLARIS in developing human heart data indeed exhibit different gene expression profiles across layers, beyond what can be attributed to differential cell type compositions. Finally, applying POLARIS to ST data from breast cancer patients, we found that POLARIS identified LDE genes reveal complex heterogeneous across-layer/region differential expression across samples and/or patients. The LDE detected genes are consistent with established knowledge regarding breast cancer pathology, including metastasis and prognosis, but offer more granular sample-level and patient-level information that can potentially empower personalized diagnosis and treatment.

Another key feature of POLARIS is its ability to leverage image data. In the ST deconvolution field, gene expression itself has proved its ability to infer cell composition. Imaging information, however, has been under-utilized. Recent work [18, 19] showed the potential of histology images accompanying ST data. We believe that histology images can be further leveraged for ST inference. For example, histology images alone are widely used to segment cells with deep learning models [13, 14]. POLARIS can take an accompanying image as input to train an image network, and employ a pre-trained image network on a completely new image. Our pre-trained POLARIS image network offers a novel method for tissue registration, which extracts and reveals tissue anatomical or functional structures either from the histological image alone or jointly with gene expression. The major barriers that prevent the full potential of integrating histological images with ST data include quality of the co-registered image and, most importantly, the absence of pathologist annotations. In order to accomplish the task with the currently available data, POLARIS incorporates training of the image network into the inference of cell composition. Instead of training a model with inferred cell composition as the goal and using MSE as the loss function, our intention is to use histology images to help with the estimation of cell composition, under the rationale that spots with similar histological images and similar neighborhoods tend to share similar cell composition. Nevertheless, we fail to demonstrate that the image improves deconvolution performance due to the limitation of the current data: single-cell level resolution ST data only provides DAPI stained images which only comprise one color panel, while spot-level ST data has no gold standard truth. Despite these limitations to quantify the performance, POLARIS with image input still achieved high accuracy among the state-of-art methods in single-cell resolution ST data, and in spot-level ST data, cell type composition inferred by POLARIS agreed with single-cell level data and the expected biological layers (e.g. glutamatergic neurons in the mouse cortex and cancer epithelial cells in the breast cancer slides). POLARIS introduces a novel approach for inferring cell composition purely from histological images that has not previously been explored by ST deconvolution. We believe that the versatile ability of POLARIS to incorporate histological images to elucidate layer-specific gene expression patterns will empower novel discoveries in spatial biology.

## Data and code availability

The processed developing human heart scRNA-seq, ISS ST data, mouse cortex SSp ST data, SSp scRNA-seq and VISp scRNA-seq reference could be obtained from the GITHUB repository of the review paper [6]: https://github.com/JiawenChenn/St-review/. The STARmap data are obtained from https://www.starmapresources.org/data and the breast cancer data are obtained from https://doi.org/10.5281/zenodo.4751624 (ST) and https://singlecell.broadinstitute.org/single_cell/study/SCP1039 (scRNA-seq reference). The processed STARmap data and breast cancer data and the source code of POLARIS could be found at GITHUB repository: https://github.com/JiawenChenn/POLARIS.

