## Supplementary information for "Cell composition inference and identification of layer-specific transcriptional profiles with POLARIS"

**Online methods**

**POLARIS inference algorithm**

We use *g* for gene index, *c* for cell index, *z* for cell type index, and *L* for layer index. $Z_c$ is the cell type of cell *c*, $L_c$ is the layer of cell *c* and $L_s$ is the layer of spot *s*. We assume that in scRNA-seq reference dataset, $X_{cg}$, expression count of gene *g* in cell *c* follows the following negative binomial distribution:

$$X_{cg} \sim NB\left(S_c\, Softplus\left(\theta_{gz_c} + T_{L_c g}\right), P_g\right)$$

$Softplus(x) = \log(1 + \exp(x)))$ is used to guarantee that $R_{gz_c} > 0$. $P_g = Sigmoid\left(o_g\right)$, where $Sigmoid(x) = \frac{1}{1+\exp(-x)}$ is the sigmoid function. $S_c$ is the library size of cell *c* and $Z_c$ is the cell type of cell *c*. $NB$ is the negative binomial distribution. $\theta_{gz_c}$ is the mean location parameter shared across layers. To capture the gene expression variation between layers, we introduce a new parameter $T_{Lg}$, a mean shift parameter for layer *L* and gene *g*. We assume that $T_{Lg} \sim N(0,1)$. However, in real world, we do not have the layer information in the scRNA-seq reference data. We make the following assumption

$$X_{cg} \sim NB\left(S_c\, Softplus\left(\theta_{gz_c}\right), P_g\right)$$

when inferring the parameters in the reference data. Estimates for the parameters are then obtained by finding the MLE (maximum likelihood estimates), given the provided scRNA-seq reference data via the gradient-based optimization using the PyTorch library in Python.

In the ST data, we make similar distributional assumption. Specifically, we assume that $X_{scg}$, expression count of gene *g* in cell *c* in spot *s* follows the following negative binomial distribution:

$$X_{scg} \sim NB\left(\beta_g Softplus\left(\theta_{gz_c} + T_{L_c g}\right), P_g\right)$$

$\beta_g$ is the parameter measuring technical bias between ST and scRNA-seq reference for gene *g*. Note that the bias parameter is gene-specific. Considering the additive property of negative binomial distribution and summing across all single cells within spot *s*, the resulting distribution of gene *g* in spot *t* also follows a negative binomial distribution:

$$X_{sg} \sim NB\left(\sum_{z=1}^{Z} \beta_g Softplus\left(\theta_{gz} + T_{L_s g}\right) n_{sz}, P_g\right)$$

*Z* is total number of cell types and $n_{sz}$ is the number of cells of type *z* in spot *s*. All the parameter are inferred using maximum a posteriori estimation (MAP) with $\widehat{\theta_g}, \widehat{P_g}$ obtained from scRNA-seq reference. At last, cell type proportion can be calculated as $v_{sz} = \frac{n_{sz}}{\sum n_{sz}}$.

**Image feature extraction using masked autoencoder (MAE)**

The MAE codes are obtained from https://github.com/facebookresearch/mae . We only use the pretrained encoder part to extract the image features [32]. We used mae_visualize_vit_large_ganloss.pth as the pretrained model. We used MAE to extract image features for the spot image and spot neighborhood image. The spot image is a $r * r$ square covering the spot. $r$ is defined specific to the dataset used (details about the used *r* could be found in the source code). The neighborhood image is defined as a $3r * 3r$ square sharing the same center point as the spot image. The two MAE-extracted 1024-length vectors are combined as the image input of the spot. The POLARIS image network combines two levels of fully connected layers. The $\theta_{gz}$ and $T_{L_s g}$ are fixed during the training of image network.

**Simulation**

We begin by generating a single-cell reference. A total of 100 genes are simulated in six cell types. The number of cells in each cell type is simulated from $N(500, 100^2)$ and rounded to the nearest integer. For each cell, we simulate the layer from $Binomial(0.3)$ where we treat a simulated value of 0 as layer 1 and value of 1 as layer 2. We then simulate the gene expression of gene *g* in cell *c* from $NB(Softplus(\theta_{gz_c} + T_{L_c g}), P_g)$, where NB denotes the negative binomial distribution. $Z_c$ represents the cell type of cell *c* and $L_c$ represents the layer of cell *c*. Both $\theta_{gz_c}$ and $T_{Lg}$ are simulated from $N(0,1)$ and $P_g$ is simulated from $Uniform(0.2,0.8)$. Note $T_{Lg}$ represents the layer-specific gene expression profile, thus allowing expression profiles to vary across layers, even for the same gene *g* and cell type. We then construct pseudo spots by randomly selecting cells in layer 1 and layer 2 with replacement from single cells simulated above. In each spot, the number of cells is determined by sampling a random number from Uniform(10,16). We simulated 50 spots in layer1 and 150 spots in layer2. For simulation with different proportion of genes with layer-specific expression profiles, we define the proportion as $m$. Then the $T_{Lg}$ for $100 * (1 - m)$ % genes are set as 0 while the $T_{Lg}$ for the rest $100 * m$ % genes are simulated from $N(0,1)$.

**Calculate between-layer fold change**

To identify LDE genes, we compare the gene expression profile across layers by comparing layer-specific location parameters:

$$Softplus(\theta_{gz} + T_{Lg})$$

Specifically, the fold change of gene *g* in layer $L$ compare to other layers in cell type *z* are calculated as

$$F_{Lg} = (\theta_{gz} + T_{Lg}) / (\sum_{L' \neq L} (\theta_{gz} + T_{L'g}) / (S - 1))$$

Where *S* is the total number of layers. Then we take the maximum fold change across cell types to quantify the across-layer fold change of gene *g*.

**LDE gene identification**

We perform permutation test to identify LDE genes. With a given layer annotation $Layer^{obs}$, we perform cell type deconvolution and infer $T_{Lg}^{obs}$ and $F_{Lg}^{obs}$. Then we randomize the layer annotation for $N$ times and generate pseudo layer annotation $Layer^{sim1}, \ldots, Layer^{simN}$. Consequently, we are able to infer $T_{Lg}^{sim1}, \ldots, T_{Lg}^{simN}$. Then the p value of gene *g* in layer $L$ is defined as

$$p\ value = \frac{\#\ |T_{Lg}^{sim}| > |T_{Lg}^{obs}|}{N}$$

Genes with $p\ value < \frac{0.05}{Total\ number\ of\ genes}$ (Bonferroni correction) and $\left|\log_2 F_{Lg}^{obs}\right| > 1$ are considered as the LDE genes in layer $L$ (marked as pink in the figures). We used $N = 10000$ (permutation times) in all the analysis.

**Data preprocessing**

For the STARmap data, we only keep genes that are present in at least 2% of spots.
In the breast cancer analysis, we only keep HER2+ subtype ST data for deconvolution evaluation. Similarly, we keep only the HER2+ patients' scRNA-seq data as reference. For scRNA-seq reference, genes expressed in at least 3 cells and cells expressing at least 200 genes are kept. Similarly for ST data, genes expressed in at least 3 spots and spots expressing at least 100 genes are kept. Only genes that exist in both scRNA-seq and ST data are employed in further analysis. Top 2000 highly variable genes (HVGs) are used. The gene subsettings are accomplished using the R package Seurat [54]. HVGs are selected using feature variance calculated by the FindVariableFeatures function with default settings.

For the heart ISS DAPI-stained image, we reverse the color and enhance the contrast using ImageEnhance function in the Pillow package [55].

**Comparing to other state-of-art methods**

We compared the performance of POLARIS with several state-of-the-art deconvolution methods developed for ST data. We followed the instructions of each method on their corresponding website. Among all the methods, only RCTD has a built-in gene filtering method, where only genes with normalized gene expression >= 0.0002 are included, and it selects cell type marker genes based on a log-fold-change threshold of 0.75 [28]. We used the default parameters of RCTD and ran RCTD in full mode. Only selected cell type marker genes were fed into RCTD. For all other methods, we used all genes without any further filtering from the preprocessed data described in the previous section.

**Tissue detection**

To use POLARIS-trained image network, we need to automatically detect the tissue section from a histological image. We utilized the filter_entropy function in https://github.com/CODAIT/deep-histopath and followed the instructions in https://developer.ibm.com/articles/an-automatic-method-to-identify-tissues-from-big-whole-slide-images-pt3/ . We used entropy, which measures tissue complexity, to detect the percentage of tissue in each spot. Areas such as the slide background are less complex than the tissue area. We used the default threshold of 5 in the analysis. Pixels with an entropy value greater than 5 are counted as tissue regions. Spots with greater than 5% tissue area is kept in the super-resolution composition inference.

**Acknowledgements**

We thank Li lab members for providing advice on data preprocessing, method selection and feedback on the manuscript. Figure 1 is created via bioRender. The research is supported by National Institutes of Health (NIH) grants (R01HL163972 and U01HG011720).

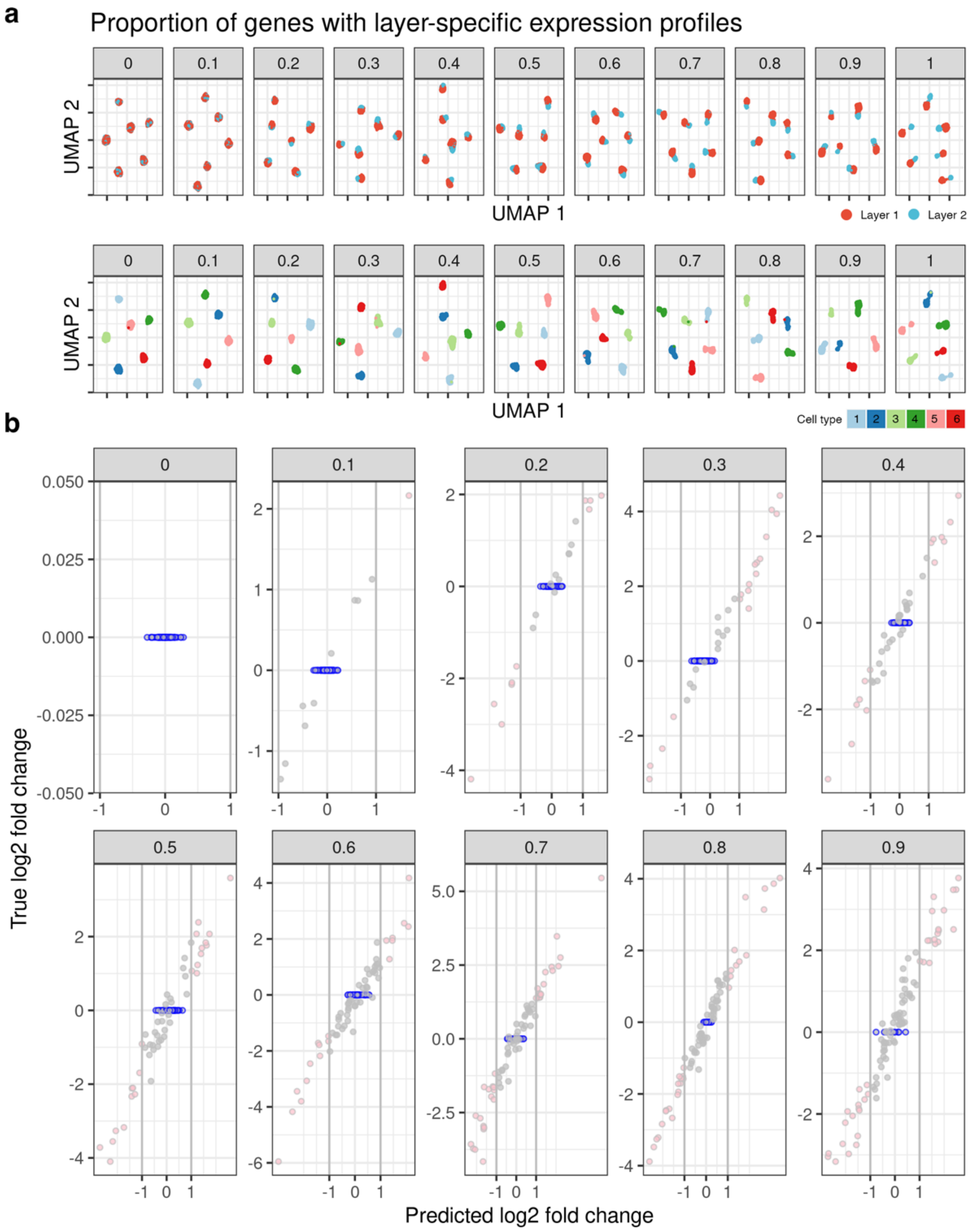


**Extended Data Fig. 1 | POLARIS inferred log2 fold change of gene expression across layers agrees with the true log2 fold change and well-controlled type-I error in simulations.** (a) UMAP visualization of cells colored by layer (top) and cell type (bottom) (b) Points with blue borders are the genes with invariant gene expression profiles across layers. Pink points mark LDE genes identified by POLARIS.

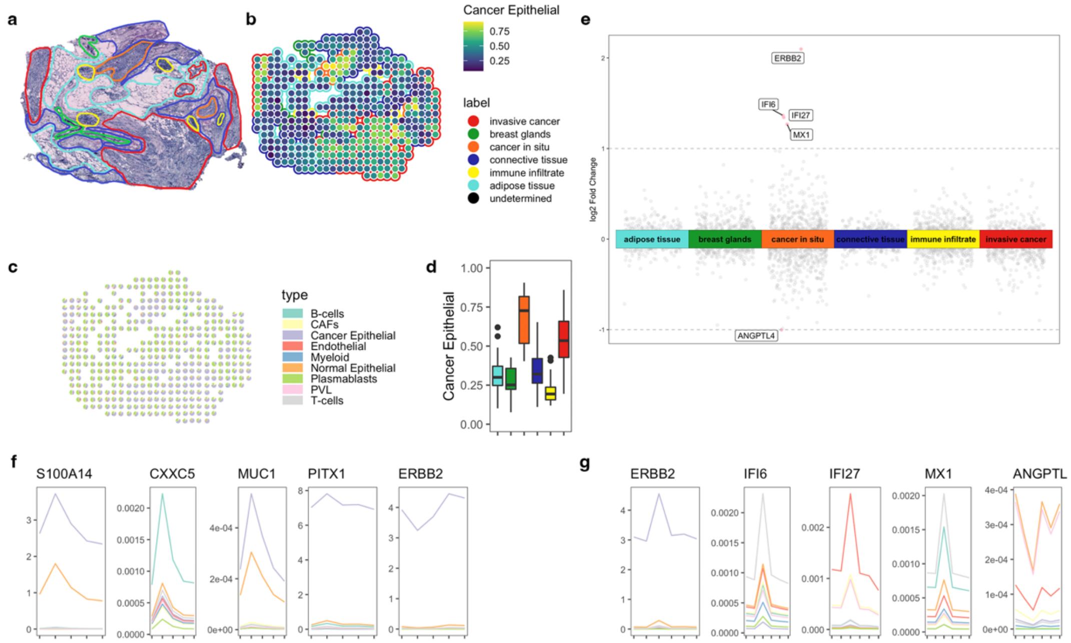

**Extended Data Fig. 2 | POLARIS inference on breast cancer slide G2.** (a) Pathologist annotation on slide G2 (b) POLARIS inferred cancer epithelial cell proportion (c) POLARIS inferred cell composition (d) Distribution of POLARIS inferred cancer epithelial cell proportions, in each layer (color scheme is the same as in a and b above) (e) POLARIS inferred log2 fold change of gene expression across layers. Points with absolute value greater than 1 are colored as pink, otherwise, gray. (f) POLARIS inferred gene expression location parameter on slide A1 (Fig. 3f-k) (g) POLARIS inferred gene expression location parameter on this slide G2.

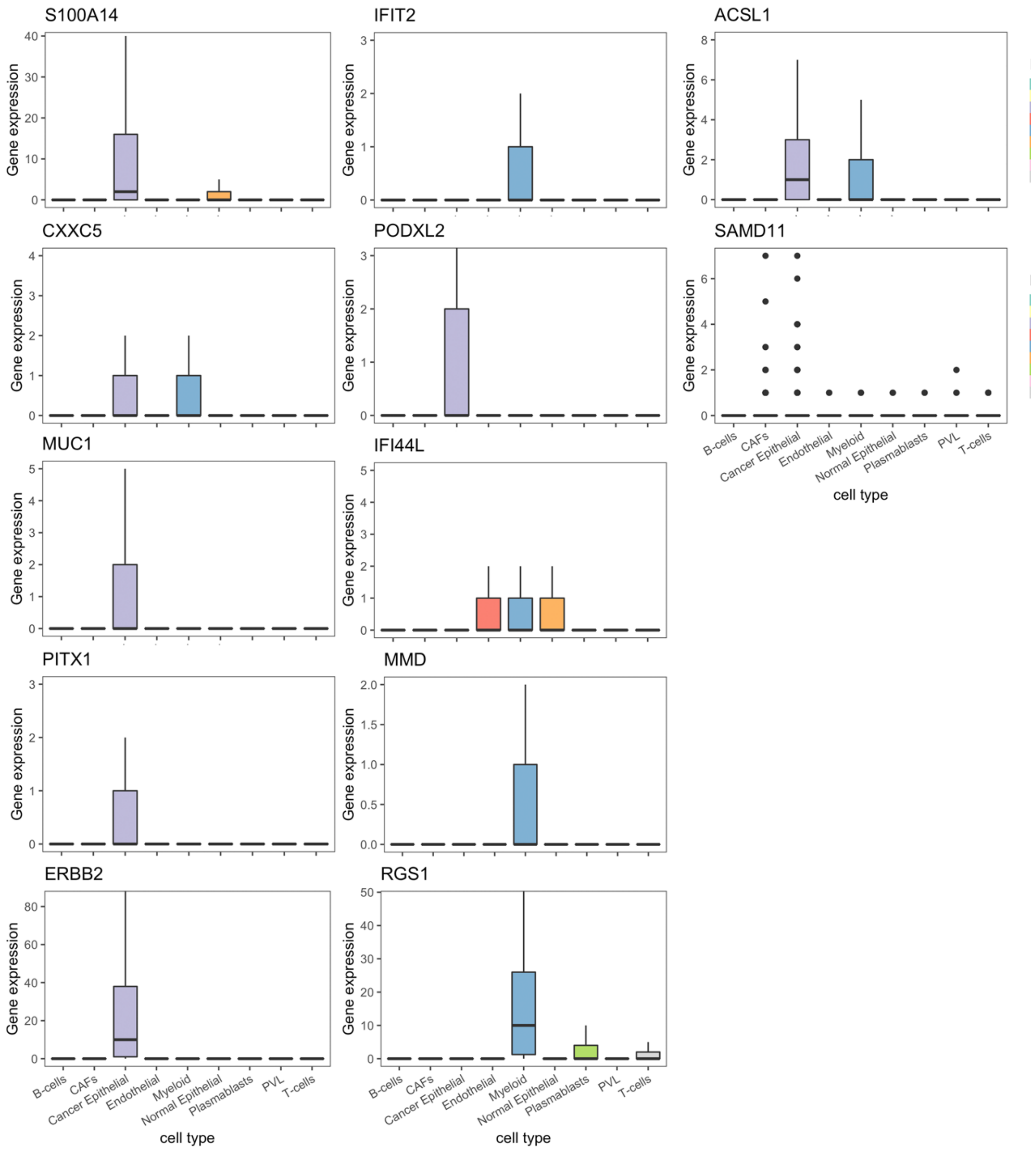


**Extended Data Fig. 3 | Cell type specific gene expression distribution of POLARIS identified LDE genes in breast cancer, based on scRNA-seq data.** The side-by-side boxes are colored by cell types.

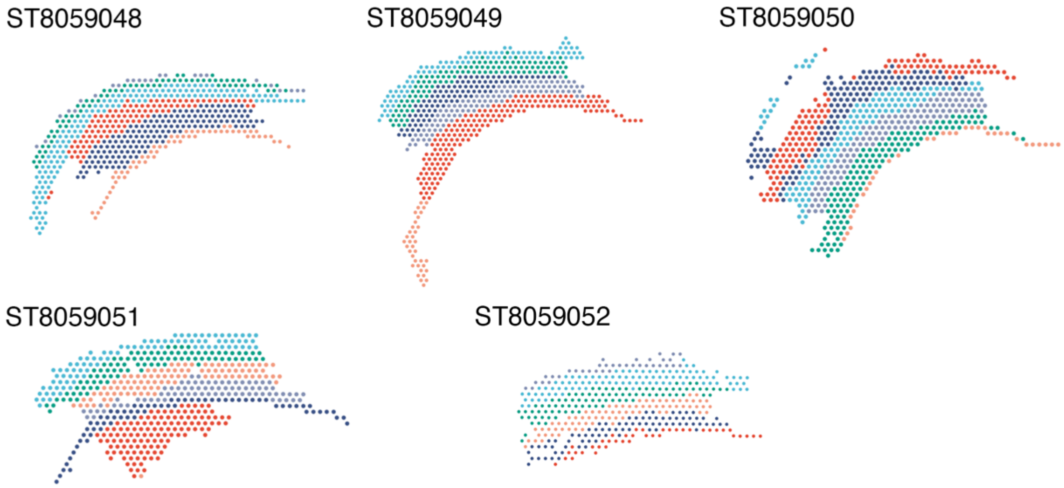


**Extended Data Fig. 4 | BayesSpace inferred clusters in the mouse cortex SSp 10x Visium data.**

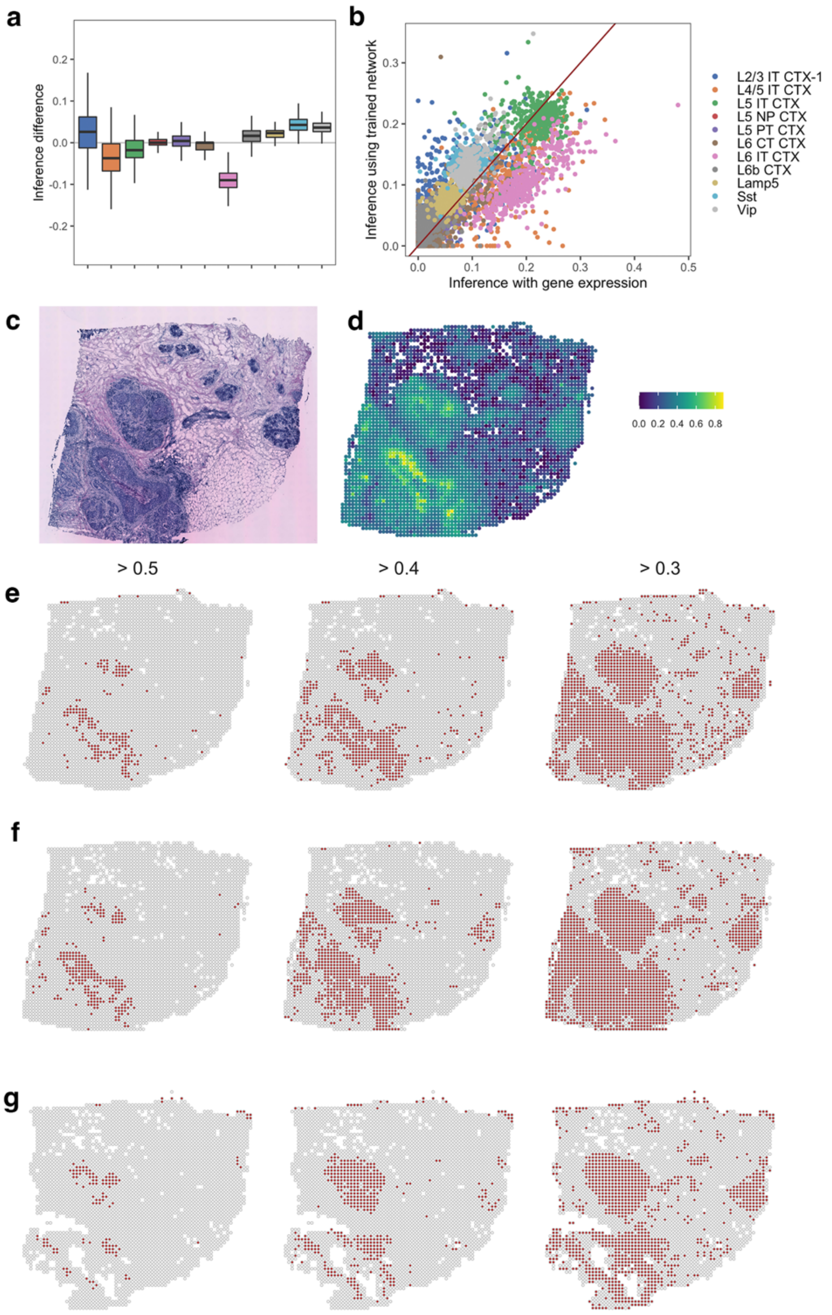


**Extended Data Fig. 5 | Cell composition inferred using POLARIS-trained image network is comparable with inference with gene expression and can be used as a new method for tissue segmentation.** (a) Difference in the inferred cell composition, between estimates using POLARIS-trained image network and estimates using gene expression, former minus latter. Each boxplot represents one cell type. (b) Estimated cell type proportions from POLARIS-trained image network and from gene expression data are correlated (for example, Peason's correlation in L2/3 IT CTX-1 : 0.52, L4/5 IT CTX : 0.75, L5 IT CTX : 0.21, L5 NP CTX : 0.29, L5 PT CTX : 0.32, L6 CT CTX : 0.76, L6 IT CTX : 0.78, L6b CTX : 0.80, Lamp5 : 0.36, Sst : 0.31, Vip : 0.69). (c) Histology image of HER2+ breast cancer slide H2 (d) Super-resolution inference of H2 using POLARIS image network trained on H1. (e-g) Tissue registration using different cancer epithelial cell proportion (>0.5, >0.4, >0.3) on slide (e) H1 (f) H2 (g) H3. Spots colored in red have a larger proportion of cancer epithelial cells than the corresponding threshold (0.5, 0.4, and 0.3 from left to right). These spots can be treated as falling in cancerous areas.
